# Dynamic Phosphorylation Regulates Eukaryotic Translation Initiation Factor 4A Activity During the Cell Cycle

**DOI:** 10.1101/2022.08.14.503931

**Authors:** Ansuman Sahoo, Robert A. Zollo, Shichen Shen, Marium Ashraf, Samantha Nelson, Gerald Koudelka, Jun Qu, Joseph Barbi, Sarah E. Walker

## Abstract

The eukaryotic translation initiation factor 4A (eIF4A) resolves mRNA structures to support protein synthesis, yet little is known about its regulation. Here we analyzed eIF4A phosphorylation during alternate stages of the cell cycle, and found three residues near the DEAD box motif (T73, T146, and S177) underwent substantial phosphorylation changes. Phosphomimetic mutations T73D and T146D led to G2/M phase arrest, and abolished eIF4A interaction with RNA, suggesting eIF4A activity is needed for completion of cell division. In addition to these repressive events, we found that S177, a site immediately adjacent to the DEAD-box, showed diametrically opposed phosphorylation, with only phosphorylated S177 present during G1/S arrest and dephosphorylated S177 peptides during G2/M arrest. Phosphomimetic S177D eIF4A increased polysome levels and enhanced normally reduced eIF4A-eIF4G-interaction during G2/M, while phosphodeficient S177A decreased polysome levels and reduced growth, suggesting phosphorylation of S177 enhances eIF4A-mediated translation during G1/S. Together these results suggest that dynamic phosphorylation of eIF4A S177 serves to stimulate translation during G1/S, while inhibitory phosphorylation of additional sites holds the potential to rapidly transition eIF4A to an inactive state and turn off translation. These results also suggest an important role for eIF4A in coupling translation to cell cycle stages.

## INTRODUCTION

Eukaryotic translation initiation begins with the recruitment of a preinitiation complex (a 40S ribosomal subunit with various eIFs and initiator tRNA) bound to the m^7^G cap at the 5’ end of the mRNA, an event mediated by the eIF4F complex, eIF4B and eIF3 (1). eIF3 binds to tRNA-containing ternary complex as part of a multifactor complex that binds the pre initiation complex (PIC), promoting a scanning-competent open conformation of the 40S subunit. eIF4F is a trimeric complex consisting of the RNA-binding scaffolding protein eIF4G, cap-binding protein eIF4E, and RNA helicase eIF4A. eIF4B is a ribosome- and RNA-binding protein that interacts with eIF4A to enhance its activity within the eIF4F complex. The activity of eIF4F and eIF4B, as well as additional helicases for highly structured mRNAs, are needed to melt structure in the 5’UTR of mRNAs and potentially to bridge the 43S complex and mRNA (2-5).

eIF4A is an ∼45 kDa highly-conserved DEAD-box protein critical for loading and likely scanning of mRNAs during translation initiation. As such, it contains two RecA domains tethered by a linker, and can occupy both an open conformation, observed in the absence of RNA, eIF4B and eIF4G, where the two domains are far apart, and partially and fully-closed conformations, in which the RecA domains move closer together to bind RNA. Importantly, the RNA- and ATP-dependent RNA helicase activity relies on switching between these open and closed conformations (6-8). Although eIF4A has weak unwinding activity on its own, eIF4G and eIF4B stimulate eIF4A by both promoting a fully closed conformation that enhances RNA binding affinity, and lengthening the time spent in the fully closed conformation for effectively coupling ATPase to unwinding (9, 10). Interaction of eIF4G with eIF4A additionally provides specificity towards single-stranded 5’ overhangs, thus biasing the unwinding activity of the eIF4F complex to 5’ untranslated regions (11). eIF4A is a highly abundant protein, and depletion confers linear decreases in translation rate, consistent with the idea that eIF4A functions through widespread melting of small RNA duplexes, rather than acting as a translocating helicase (12). A recent cryo-EM structure suggests that eIF4A, as part of the eIF4F complex, resides at the exit channel of the 40S subunit, where it is thought to resolve mRNA structure near the cap and subsequently loop mRNA out of the complex via successive ATPase activity (13). Consistent with a role for eIF4A in both PIC loading and scanning, in vitro experiments following kinetics of ribosome loading, and ribosome profiling data in yeast suggest that while all mRNAs show some dependence on eIF4A for translation, those mRNAs most affected by its inactivation (i.e., eIF4A-hyperdependent transcripts) have longer than average, and highly structured 5’UTRs and open reading frames (4, 14).

Several large-scale mass spectrometry studies reported phosphorylated peptides for eIF4A, and a number of the reported phosphosites reside in close proximity to the conserved DEAD motif and RNA-binding region of the protein (15, 16). Phosphorylation of one such site, T164 (the equivalent of T146 in baker’s yeast eIF4A), was previously analyzed in *Arabidopsis thaliana* and shown to accumulate during cell cycle arrest (G2/M) by nocodazole. The phosphorylated Threonine resides in a consensus Cdc28/CDK1/CDKA site in a region abutting the DEAD-box helix, and CDKA both binds to and mediates phosphorylation of the plant factor at this site during mitosis. Replacement of T164 with a charged Aspartic Acid residue led to growth inhibition, diminished fertility, and abolished the activity of wheat germ factor in translation in vitro, consistent with inhibition of eIF4A and translation upon phosphorylation of this residue (17, 18). Peptides indicating phosphorylation of the equivalent residue have been identified in HeLa and K562 cell lines (19) in addition to the yeast factor, and the highly conserved region surrounding this residue encompass a CDK1/Cdc28 site, suggesting phosphorylation of this residue could serve a role across eukaryotes in modulating eIF4A activity to control translation during cell cycle transitions. However, it is unclear whether additional phosphorylated sites also affect translation, and the degree to which eIF4A function is modulated in the cell cycle in other eukaryotes, warranting further study.

To further explore the function of T146 and other phosphosites of eIF4A, we analyzed the degree of phosphorylation, and effects of preventing or mimicking phosphorylation of conserved phosphosites of eIF4A using *Saccharomyces cerevisiae* and purified yeast proteins. We found that Aspartic acid substitutions of two conserved sites near the DEAD-box motif, T73 and T146, were inhibitory for eIF4A RNA binding and growth in yeast. Phosphorylation of T146, was heightened during mitotic arrest relative to G1/S phase arrest, consistent with previous findings for this residue in plants. Phosphomimetic T to D substitutions of either T73 or T146 led to mitotic arrest, but only in the absence of WT eIF4A, suggesting eIF4A activity is needed at the late M to G1 transition, and inhibiting this activity could block synthesis of a protein necessary to complete division. Surprisingly, we also found a conserved residue, S177, immediately adjacent to the DEAD box that was completely phosphorylated during G1/S and dephosphorylated during G2/M arrest. Inhibiting phosphorylation of S177 by Alanine substitution decreased the number of polysomes present in asynchronously growing cells, suggesting this site stimulates eIF4A activity during translation initiation. Together, these data suggest that eIF4A activity is capable of both rapid shutdown as well as stimulatory fine-tuning through distinct phosphorylation events. Besides describing a conserved but understudied mechanism for the regulation of eIF4A activity, this study highlights the importance of post-translational eIF4 factor modifications in coupling translation to a fundamental cellular process (i.e., cell cycle progression) and the possibility for additional controls of eIF4A activity in response to myriad cellular changes.

## MATERIAL AND METHODS

### Construction of yeast strains and plasmids

All strains, plasmids, and primers used in this study are listed in Tables 1, 2, and 3, respectively. For construction of pAS1 and pAS3, a *TIF1* allele containing the native promoter and terminator was amplified from BY4741 genomic DNA and Gibson-assembled with BamHI cut pRS416 and pRS415 backbones. Phosphorylation site mutations and gRNA sequences were introduced into vectors by PCR amplification with mutagenic primers (Table 3) with Phusion (Thermo) or PfuUltra II (Stratagene) polymerase, followed by transformation of DpnI-digested products. For simultaneously mutating both copies of the eIF4A gene within the yeast genome, a gRNA was identified for targeting the region near T146 of *TIF1* and *TIF2* using the CRISPR direct website, and then cloned into the pML104 vector harboring *S. pyogenes* Cas9 by PCR-mutagenesis to generate a new targeting vector, pMD2. This vector and preannealed homology-directed repair oligos were transformed into BY4741 to generate YSW15 and 16. Resulting mutations in each gene were verified by sequencing allele*-*specific colony PCR products. For construction of strain YSW30, the *TIF1* and *TIF2* genes encoding eIF4A were deleted using the same CRISPR/Cas9 and homology directed repair strategy (20, 21). BY4741 cells were transformed with the pCAS-derived KanMX vector (pAS2) and two double-stranded DNAs for homology-directed repair that fused the 5’ and 3’UTRs of the *TIF1* and *TIF2* genes. The transformation also contained a single-copy URA vector (pAS1) encoding a *TIF1* gene with silent mutations in the gRNA annealing site that conferred resistance to Cas9-targeting without affecting growth. Isolates were subsequently grown on SC-Ura without G418 and screened for loss of the pAS2 vector by patching onto G418-containing media, then verified to have lost both chromosomal eIF4A genes by production of distinct colony PCR products.

**Table 1:** Yeast strains used in this study.

| **Strain Name** | **Genotype** |
| --- | --- |
| YSW30 | *MATa, his3Δ1, leu2Δ0, ura3Δ0, met15Δ0, tif1Δ0/tif2Δ0 [pAS1]* |
| YSW47 | *MATa, his3Δ1, leu2Δ0, ura3Δ0, met15Δ0, tif1Δ0/tif2Δ0 [pAS1][pAS3]* |
| YSW48 | *MATa, his3Δ1, leu2Δ0, ura3Δ0, met15Δ0, tif1Δ0/tif2Δ0 [pAS1][pAS4]* |
| YSW49 | *MATa, his3Δ1, leu2Δ0, ura3Δ0, met15Δ0, tif1Δ0/tif2Δ0 [pAS1][pAS5]* |
| YSW50 | *MATa, his3Δ1, leu2Δ0, ura3Δ0, met15Δ0, tif1Δ0/tif2Δ0 [pAS1][pAS6]* |
| YSW51 | *MATa, his3Δ1, leu2Δ0, ura3Δ0, met15Δ0, tif1Δ0/tif2Δ0 [pAS1][pAS7]* |
| YSW52 | *MATa, his3Δ1, leu2Δ0, ura3Δ0, met15Δ0, tif1Δ0/tif2Δ0 [pAS1][pRS415]* |
| YSW111 | *MATa, his3Δ1, leu2Δ0, ura3Δ0, met15Δ0, tif1Δ0/tif2Δ0 [pAS1][pAS10]* |
| YSW124 | *MATa, his3Δ1, leu2Δ0, ura3Δ0, met15Δ0, tif1Δ0/tif2Δ0 [pAS3]* |
| YSW125 | *MATa, his3Δ1, leu2Δ0, ura3Δ0, met15Δ0, tif1Δ0/tif2Δ0 [pAS57]* |
| YSW126 | *MATa, his3Δ1, leu2Δ0, ura3Δ0, met15Δ0, tif1Δ0/tif2Δ0 [pAS58]* |
| YSW246 | *MATa, his3Δ1, leu2Δ0, ura3Δ0, met15Δ0, tif1Δ0/tif2Δ0 [pAS1][pAS25]* |
| YSW247 | *MATa, his3Δ1, leu2Δ0, ura3Δ0, met15Δ0, tif1Δ0/tif2Δ0 [pAS1][pAS26]* |
| YSW248 | *MATa, his3Δ1, leu2Δ0, ura3Δ0, met15Δ0, tif1Δ0/tif2Δ0 [pAS1][pAS27]* |
| YSW249 | *MATa, his3Δ1, leu2Δ0, ura3Δ0, met15Δ0, tif1Δ0/tif2Δ0 [pAS1][pAS28]* |
| YSW250 | *MATa, his3Δ1, leu2Δ0, ura3Δ0, met15Δ0, tif1Δ0/tif2Δ0 [pAS1][pAS29]* |

**Table 2:** Plasmids used in this study.

| Addgene ID | Plasmid Name | Relevant Gene(s) | Markers | Parental Vector | Template | Primers |
| --- | --- | --- | --- | --- | --- | --- |
| TBD | pMD2 | *TIF1/2* | *E. coli* AmpR,<br>Yeast *URA3* | pML104 | N/A | MB26/MB27 |
| TBD | pAS1 | *TIF1* | *E. coli* AmpR,<br>Yeast *URA3* | pRS416 | BY4741<br>DNA | AS1/2/3/4 |
| TBD | pAS2 | *Cas9, gRNA to TIF1/2* | *E. coli* AmpR,<br>Yeast *KANMX* | pCAS | N/A | AS5/6 |
| TBD | pAS3 | *TIF1* | *E. coli* AmpR,<br>Yeast *LEU2* | pRS415 | BY4741<br>DNA | AS1/4 |
| TBD | pAS4 | *tif1*<sup>^S2A</sup> | *E. coli* AmpR,<br>Yeast *LEU2* | pAS3 | N/A | MB18/19 |
| TBD | pAS5 | *tif1*<sup>^S2D</sup> | *E. coli* AmpR,<br>Yeast *LEU2* | pAS3 | N/A | MB22/23 |
| TBD | pAS6 | *tif1*<sup>^T146A</sup> | *E. coli* AmpR,<br>Yeast *LEU2* | pAS3 | N/A | SW279/280 |
| TBD | pAS7 | *tif1*<sup>^T146D</sup> | *E. coli* AmpR,<br>Yeast *LEU2* | pAS3 | N/A | SW281/282 |
| TBD | pAS10 | *tif1*<sup>^T146E</sup> | *E. coli* AmpR,<br>Yeast *LEU2* | pAS3 | N/A | SW283/284 |
| TBD | pAS25 | *tif1*<sup>^T73D</sup> | *E. coli* AmpR,<br>Yeast *LEU2* | pAS3 | N/A | AS66/67 |
| TBD | pAS26 | *tif1*<sup>^S68A</sup> | *E. coli* AmpR,<br>Yeast *LEU2* | pAS3 | N/A | AS74/75 |
| TBD | pAS27 | *tif1*<sup>^S68D</sup> | *E. coli* AmpR,<br>Yeast *LEU2* | pAS3 | N/A | AS76/77 |
| TBD | pAS28 | *tif1*<sup>^S129A</sup> | *E. coli* AmpR,<br>Yeast *LEU2* | pAS3 | N/A | AS78/79 |
| TBD | pAS29 | *tif1*<sup>^S129D</sup> | *E. coli* AmpR,<br>Yeast *LEU2* | pAS3 | N/A | AS80/81 |
| TBD | pAS31 | *tif1*<sup>^T73A</sup> | *E. coli* AmpR,<br>Yeast *LEU2* | pAS3 | N/A | AS118/119 |
| TBD | pAS57 | *tif1*<sup>^S177A</sup> | *E. coli* AmpR,<br>Yeast *LEU2* | pAS3 | N/A | SW377/378 |
| TBD | pAS58 | *tif1*<sup>^S177D</sup> | *E. coli* AmpR,<br>Yeast *LEU2* | pAS3 | N/A | SW379/380 |

TBD, To be determined; Addgene IDs for each plasmid will be added prior to publication.

**Table 3:**
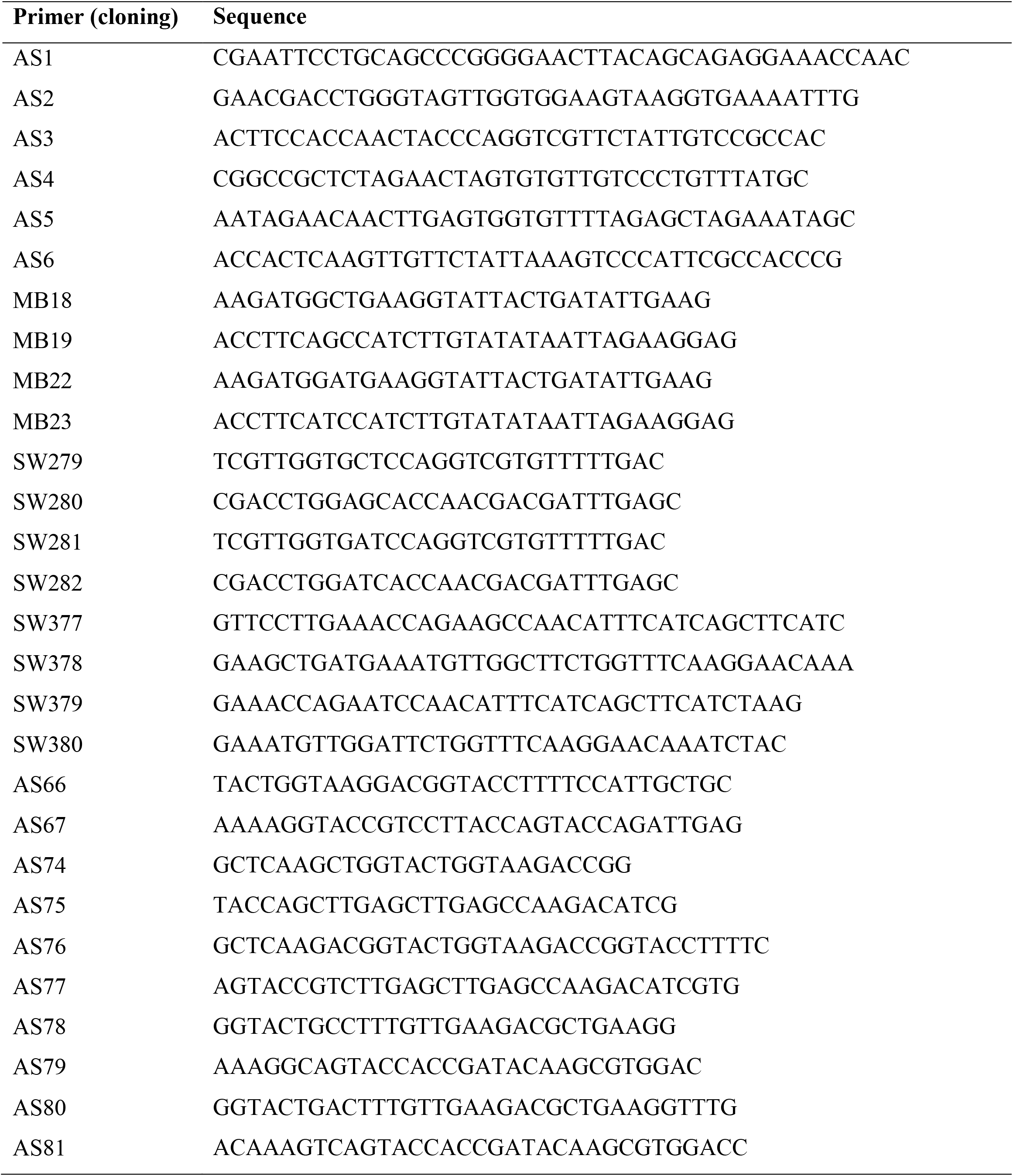
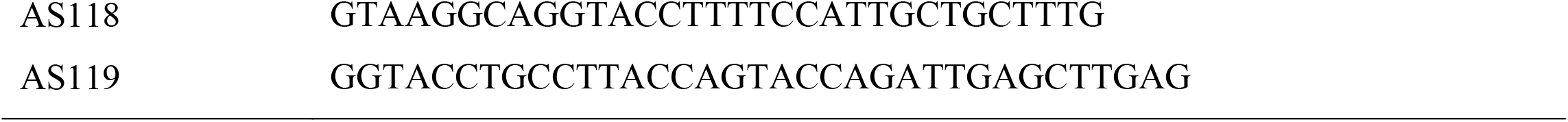
Primers used in this study.

### Yeast growth

Yeast strains were cultivated in liquid or on solid (2% agar) YPD media (Teknova, USA) or Synthetic Complete (SC) media (20 g/L glucose, 1.71 g/L yeast nitrogen base without amino acids, SC dropout mix, 5 g/L ammonium sulfate; Sunrise Scientific Products, USA) as indicated, at 30°C. Serial 10-fold dilutions (starting with OD_600_ of 1 from a fresh overnight culture) were plated on selective media and grown at 30°C for spotting assays. To arrest cells, BY4741 yeast cells were grown in 50 mL YPD media to an OD_600_ between 0.3-0.4 before adding arresting reagent. For G2/M arrest, nocodazole was added to a final concentration of 15 µg/mL and for G1/S-phase arrest, hydroxyurea was added to a final concentration of 200 mM.

### Antibodies

Polyclonal rabbit antibodies against untagged yeast eIF4A and eIF4G were described previously (22). Polyclonal anti-eIF4A antibody was further purified from 40 ml of serum by running over a 5 ml Protein A column (Acrobiosystems Inc., MA-0422-C5) for immunoprecipitation experiments. Custom antibodies were verified against recombinant protein (23) and optimized for linear behavior using lysates from BY4741 yeast. Antibodies were used at the following concentrations for western blotting: rabbit anti-eIF4A (1:20000 dilution, custom), rabbit anti-eIF4G (1:1000, custom), and mouse anti-HA (1:5000, Invitrogen, 26183).

### Immunoprecipitation

To detect eIF4G association with eIF4A, cells were pelleted at 0.3 OD_600_ and lysed using glass beads as described previously (24) in 75 µl of cold lysis buffer (50 mM Tris-HCl (pH 7.5), 50 mM NaCl, 0.1% Triton X-100, 10% glycerol, 1 mM EDTA, 10 ug/ml aprotinin, 10ug/ml leupeptin, 10ug/ml pepstatin, 1 mM 4-(2-aminoethyl) benzenesulfonyl fluoride (AEBSF), 1 mM DTT, Roche C0mplete protease inhibitor tablets (EDTA-free)). 14 µg of purified anti-eIF4A antibody were incubated with 50 µl of Protein A Dynabeads, washed once with 0.5 ml of 0.1 M sodium phosphate buffer (pH 8), then crosslinked by adding SPDP to a final concentration of 20 mM and incubating 1 hour at room temperature. Antibody-crosslinked beads were washed once with 0.5 ml 0.15 M glycine (pH 3.5) and twice with 0.5 ml lysis buffer before incubating clarified lysate with the beads overnight. The beads were washed three times with lysis buffer before and eluted with Laemmli buffer lacking reducing agent to prevent crosslink reversal.

### Western analysis

Lysates and immunoprecipitated samples were heated in Laemmli buffer, run on 10% SDS-PAGE gels, and transferred to PVDF membranes using the Trans-blot Turbo (Biorad). Membranes were blocked and blotted with antibody in TBST with 5% milk, and HRP-secondary antibodies were visualized using ECL reagent (BioRad). Experiments were repeated two or more times from biological replicates.

### Proteolytic digestion

An in-gel digestion protocol was adopted in the current study for analysis of eIF4A phosphosites by LC-MS. Immunoprecipitated eIF4A from cells arrested with hydroxyurea or nocodazole, as described above, was identified on an SDS PAGE gel by comparison with purified recombinant eIF4A (23) and verified to comigrate with yeast eIF4A in lysate by western blotting. Destained gel bands were first cut into small cubes (1-2 mm in each dimension) using a clean scalpel and transferred to new LoBind tubes (Eppendorf). Gel cubes were dehydrated by incubating in 500 μL acetonitrile (ACN) for 5 min with constant vortexing, and liquid was removed (all dehydration steps below followed the same procedure unless specified). After repeating the dehydration step twice, gel cubes were kept at 37°C in a thermomixer (Eppendorf) for 5 min to completely evaporate ACN. Protein was sequentially reduced by 100 μL 10 mM DTT at 56°C for 30 min and alkylated in 100 μL 25 mM IAM at 37°C in darkness for 30 min. Both steps were performed with rigorous vortexing in a thermomixer. Gel cubes were then dehydrated three times and incubated in 200 μL 0.0125 μg/μL trypsin (re-constituted in 50 mM pH 8.4 Tris-FA) on ice for 30 min. Excess trypsin was removed and replaced by 200 μL 50 mM pH 8.4 Tris-FA, and samples were incubated at 37°C overnight with constant vortexing in a thermomixer. Digestion was terminated by addition of 20 μL 5% FA and incubation for 15 min with constant vortexing, and liquid was transferred to new LoBind tubes. Gel bands were then dehydrated by sequential incubation with 500 μL 50% ACN in 50 mM Tris-FA and 500 μL ACN, each for 15 min with constant vortexing, and liquid from all steps above was combined. Protein digest was dried down in a SpeedVac and re-constituted in 40 μL 1% ACN and 0.05% trifluoracetic acid (TFA) in ddH2O with 10-min gentle vortexing. Samples were centrifuged at 20,000 g, 4 °C for 30 min, and supernatant was carefully transferred to LC vials for analysis.

### LC-MS analysis

The LC-MS system consists of a Dionex Ultimate 3000 nano LC system, a Dionex Ultimate 3000 micro LC system with a WPS-3000 autosampler, and an Orbitrap Fusion Lumos mass spectrometer. A large-inner diameter (i.d.) trapping column (300-um i.d. x 5 mm) was implemented before the separation column (75-um i.d. x 65 cm, packed with 2.5-um Xselect CSH C18 material) for high-capacity sample loading, cleanup and delivery. For each sample, 8 μL derived peptides was injected twice consecutively for LC-MS analysis. Mobile phase A and B were 0.1% FA in 2% ACN and 0.1% FA in 88% ACN. The 180-min LC gradient profile was: 4% for 3 min, 4–11% for 5 min, 11–32% B for 117 min, 32–50% B for 10 min, 50–97% B for 5 min, 97% B for 7 min, and then equilibrated to 4% for 27 min. The mass spectrometer was operated under data-dependent acquisition (DDA) mode with a maximal duty cycle of 3 s. MS1 spectra was acquired by Orbitrap (OT) under 120k resolution for ions within the m/z range of 400-1,500. Automatic Gain Control (AGC) and maximal injection time was set at 120% and 50 ms, and dynamic exclusion was set at 45 s, ± 10 ppm. Precursor ions were isolated by quadrupole using a m/z window of 1.2 Th, and were fragmented by high-energy collision dissociation (HCD). MS2 spectra was acquired by Ion trap (IT) under Rapid scan rate with a maximal injection time of 35 ms. Detailed LC-MS settings and relevant information can be found in a previous publication by Shen et al. (25).

### Data processing

LC-MS files were searched against *Saccharomyces cerevisiae* Swiss-Prot eIF4A protein sequence (Uniprot Protein Accession Number P10081) using Sequest HT embedded in Proteome Discoverer 1.4 (Thermo Fisher Scientific). Searching parameters include: 1) Precursor ion mass tolerance: 20 ppm; 2) Product ion mass tolerance: 0.8 Da; 3) Maximal missed cleavages per peptide: 2; 4) Fixed modifications: Cysteine (C) carbamidomethylation; 5) Dynamic modifications: Methionine (M) oxidation, peptide N-terminal acetylation, and Serine/Threonine/Tyrosine (S/T/Y) phosphorylation. Results were exported and manually curated by Microsoft Excel.

### Polysome profiling

Polysome analysis was performed as described previously (22). In brief, yeast strains were cultured in SC-Leu media at 30°C to an OD_600_ of 0.5. Cycloheximide (Gold biotechnology, USA) was added to a final concentration of 50 µg/ml and incubated for 5 minutes at 30°C with shaking before collecting cells by pelleting in centrifuge bottles packed with ice. Pellets were resuspended in 1/3 of the pellet weight of breaking buffer (20 mM Tris-HCl at pH 7.5, 50 mM KCl, 10 mM MgCl_2_, 1 mM DTT, 200 μg/mL heparin, 50 μg/mL cycloheximide, and 1 Complete EDTA-free Protease Inhibitor Tablet [Roche]/50 mL buffer), dropped into liquid nitrogen, and lysed in a Nitrogen Mill for 10 Cycles following a precool of 15 minutes with the following parameters: 1 minute run, 2 minutes cool, rate=15 (Spex Sample Prep, USA). 20 A_260_ units of lysates were resuspended in 1.5 volumes of the pellet weight of ice-cold breaking buffer and clarified by spinning at 14,000 rpm for 15 minutes at 4°C prior to separation by velocity sedimentation on 10-50% Sucrose gradients (20 mM Tris-HCl [pH 7.5], 50 mM KCl,10 mM MgCl2,1 mM DTT, 10-50% sucrose mixed using the 10-50% SW-41 gradient program on the BioComp gradient station) by centrifugation at 40,000 rpm for 3h at 4°C in a Beckman SW41 rotor.

### Cell cycle and size analysis by flow cytometry

Yeast cells harboring both WT and mutant copies of plasmid-encoded *TIF1* were spread plated on SC-Leu media containing 5-FOA to evict the WT copy prior to SYTOX green staining and flow cytometry (26). Cells were scraped from this plate after 24 hours and fixed with two volumes of 95% ethanol at -20°C for 24 hours, then rehydrated twice in 800 µl 50 mM sodium citrate buffer (pH 7.2). Staining was performed by addition of RNaseA (20 µg/ml) and SYTOX Green (2.5 µM) in a final volume of 300 µL sodium citrate at 37°C for 1 h. Proteinase K was then added to a final concentration of 0.4 µg/ml and incubated for an additional 1 h. at 55°C and then 4°C overnight. Stained cells were sonicated at 50% amplitude for 30 sec (Branson Sonicator) and run on an LSRII flow cytometer using a 488nm laser, a 530/30BP filter, and FACS-DIVA software (BD Biosciences) housed in the Roswell Park Comprehensive Cancer Center Flow Cytometry and Imaging Shared Resource. 488nm voltages (SYTOX Green signal) were set as to arrange the haploid (1N) DNA concentration at 20% of the overall linear axis and diploid (2N) population at 40% of the linear axis. Approximately 200,000 events were collected per sample and integrated SYTOX Green signal was analyzed using Flowjo software. Cells were gated as in past reports to remove both debris and confounding doublet events, and histograms of SYTOX Green signal and forward scatter, indicative of relative DNA content and cell size, respectively, were generated. Percentage of cells in G1, S, and G2/M phase were determined as previously described (26).

## RESULTS

### Binding of eIF4A to eIF4G, an indication of mRNP activation, was disrupted during mitosis

A universally conserved residue of eIF4A near the DEAD box was reported to be phosphorylated heavily during mitotic arrest, and mimicking phosphorylation inhibited eIF4A activity in vitro and in plants. Translation inhibition has also been reported to occur during mitosis in various organisms (27, 28), so we hypothesized that phosphorylation of eIF4A may be a conserved mechanism for translational control during specific stages of the cell cycle. In order to further study the function of phosphorylation of eIF4A as a modulator of its activity, we chose to first evaluate whether functional interactions of eIF4A are affected when arresting cells in G1/S by use of hydroxyurea, or in G2/M by use of nocodazole. We then immunoprecipitated eIF4A following arrest in the opposing phases (Figure 1A). We found by western blotting of these samples that nearly 3-fold less eIF4G was associated with eIF4A upon nocodazole arrest (“M”) than following hydroxyurea arrest (“S”), despite equivalent levels of eIF4G observed in the nocodazole-arrested cells relative to hydroxyurea. We speculated that differential modification of eIF4A and/or eIF4G could be responsible for enhanced interaction during G1/S and/or the impeded interaction and resultant translation deficits observed and reported during G2/M.

**Figure 1.**
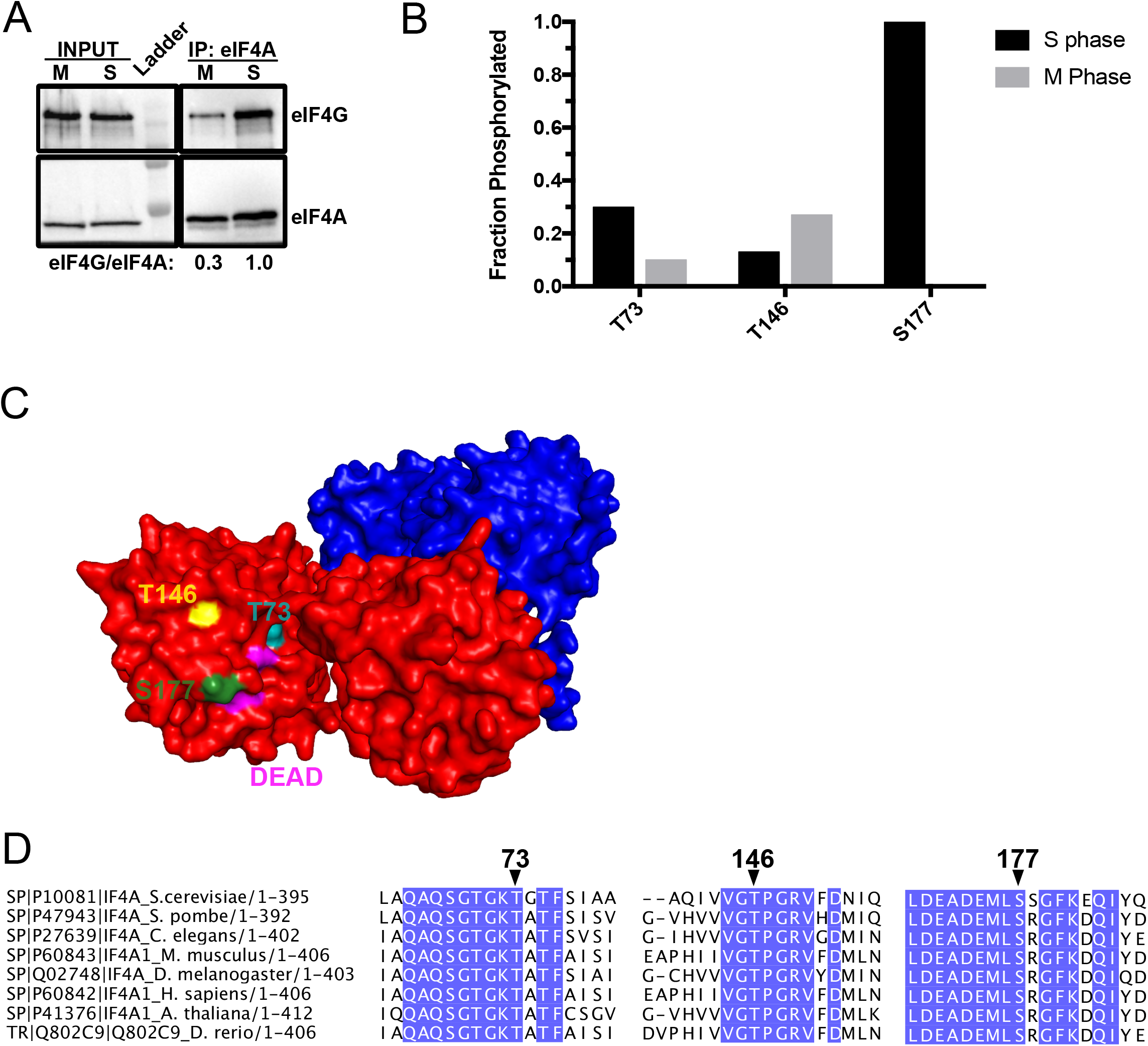
Changes in eIF4A phosphorylation state coincide with altered eIF4G interaction during mitotic arrest. **A**. eIF4G association with immunoprecipitated eIF4A following 2-hour hydroxyurea (G1/S arrest) or nocodazole (G2/M arrest) treatment of log phase yeast determined by western blot **B**. Major sites of phosphorylation that showed differences in G1/S vs G2/M arrest. **C**. Structure of eIF4A (red) bound to eIF4G (Blue) with surface-exposed residues that showed changes in phosphorylation: Threonine 73 (teal), Threonine 146 (yellow) and Serine 177 (green) around the DEAD box motif (magenta) D. Multiple sequence alignment of regions surrounding phosphorylated residues of eIF4A from model eukaryotes was generated using indicated sequences from the pdb. Blue indicates residues that showed universal conservation.

### eIF4A was differentially phosphorylated during G2/M and G1/S arrest

To further investigate the possibility that cell-cycle phase-specific phosphorylation of eIF4A affects function during translation, we next arrested cells during G1/S using hydroxyurea and G2/M using nocodazole as above, immunoprecipitated eIF4A, and performed LC/MS of the eIF4A sample from each culture to determine sites of phosphorylation. We obtained ∼90% coverage of the eIF4A protein sequence (∼700 and ∼1000 peptide spectrum matches in G1/S and G2/M samples, respectively). We then mapped the position of those phosphorylated residues within high and medium confidence peptides that were observed >5 times. We focused our investigation on those surface-exposed sites in the active site region, proximal to the DEAD box, which showed >20% phosphorylation under at least one condition, and which showed a >2-fold difference in phosphorylated peptide ratio between S and M phase, limiting our analysis to three phosphorylated residues (Figure 1B). Surprisingly, we found the heaviest phosphorylation occurred at a Serine immediately adjacent to the DEAD motif, S177, during G1/S arrest (40 phosphorylated peptides, no unphosphorylated form observed upon hydroxyurea arrest). Strikingly, zero peptides containing phosphorylated S177 were observed during G2/M arrest (6 unphosphorylated peptides), suggesting phosphorylation of this residue could control a relevant cell cycle function. Of the G1/S-specific S177 phosphorylated peptides, 20% also showed phosphorylation of S178 (8 peptides showed phosphorylation of both residues), but S178 was not observed phosphorylated independently of S177. Other residues around the active site showed less dramatic shifts as a function of cell cycle stage or a lower ratio of phosphorylated:dephosphorylated peptides. We found that the previously characterized T146 residue, located in a helix abutting the DEAD box motif (Figure 1C), showed elevated phosphorylated:unphosphorylated peptide ratio during mitosis as expected based on reports in plants (27% phosphorylated in nocodazole versus 13% in hydroxyurea arrest (18)). In contrast, T73, on the opposite side of the DEAD box motif, showed a 2.4-fold reduction in phosphorylated:dephosphorylated peptide ratio during mitotic arrest (24% phosphorylated in S phase, and 10% phosphorylated in M phase). All three residues were universally conserved in the eIF4A sequences of model eukaryotes (Figure 1D), further suggesting their potential importance in controlling eIF4A activity and cell cycle regulation across species.

### Mutations of phosphosites surrounding the DEAD box of eIF4A inhibited growth

To investigate the likely effects of phosphorylation of these sites, we first mutated each Serine or Threonine to phosphodeficient Alanine, and phosphomimetic Aspartate and/or Glutamate. We then analyzed the ability of each mutation to complement the effect of eIF4A gene deletion by plasmid shuffling in a strain lacking both chromosomal eIF4A genes, *TIF1* and *TIF2* (Figure 2). Counterselection to replace a single plasmid-encoded WT eIF4A gene (*TIF1*) on a *URA3*-containing plasmid with the same WT eIF4A allele, containing native promoter and terminator on a *LEU2*-containing plasmid produced similar levels of cell growth. Meanwhile, replacement with the corresponding empty vector control lacking the *TIF1* gene yielded no growth, indicating lack of an eIF4A gene led to lethality as expected.

**Figure 2.**
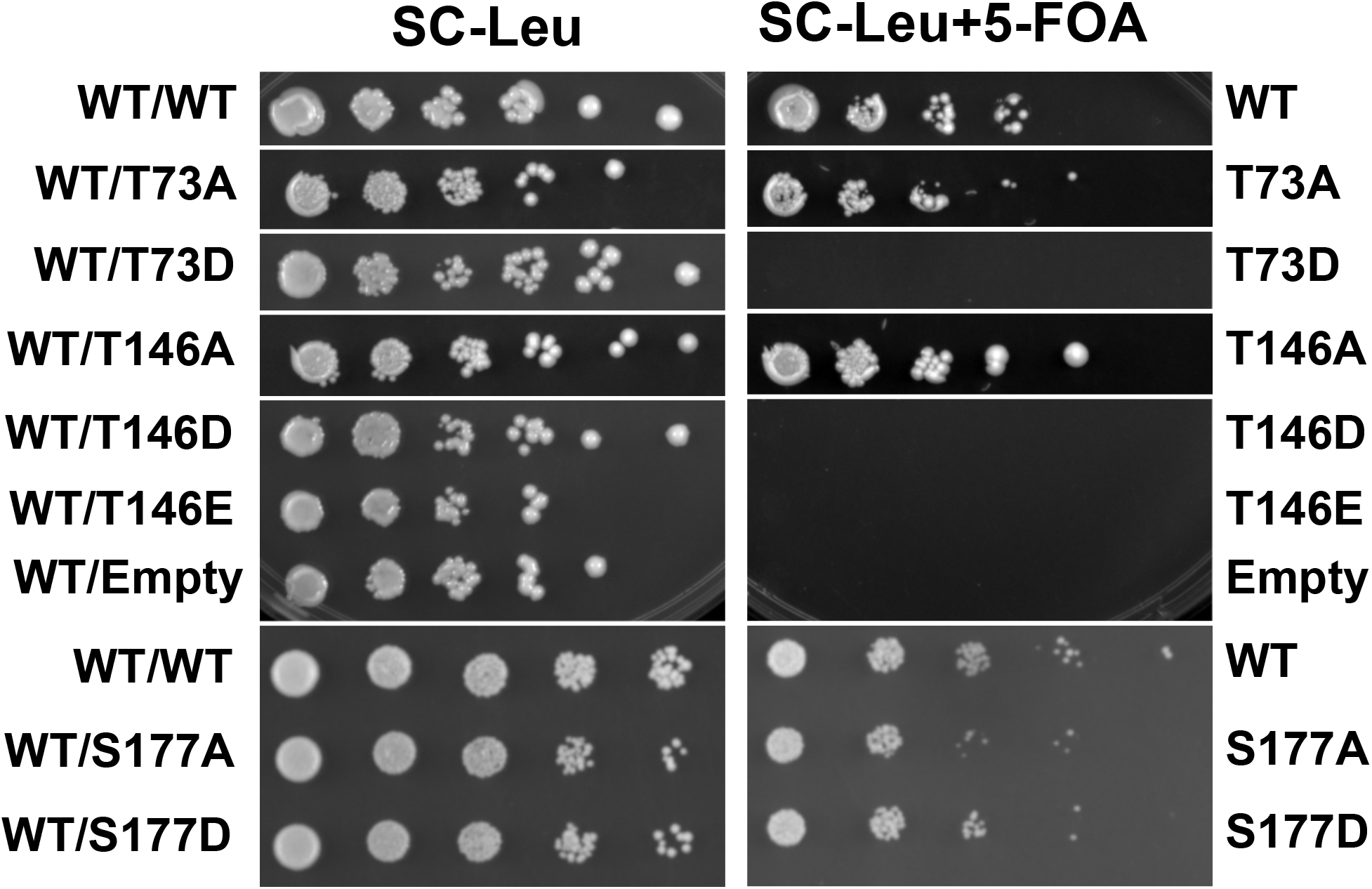
Mutations that disrupt dynamic phosphorylation of eIF4A diminish or abolish growth. Yeast *TIF1/TIF2* (encoding eIF4A) null strains harboring single copy plasmid-borne *TIF1* genes with indicated phosphorylation site mutations were spotted as 10-fold dilution series on media to allow expression with (SC-Leu) or without (SC-Leu+5-FOA) an additional WT copy. Growth was reproducibly diminished by expression of single copy eIF4A harboring phosphodeficient S177A and abolished by expression of phosphomimetic T73D, T146D and T146E. Strains were spotted 3 or more times.

We then tested whether preventing phosphorylation or mimicking constitutive phosphorylation affected growth. Plasmids harboring S177D and T146A supported growth to a similar extent as the WT *TIF1* plasmid, suggesting inability to dephosphorylate S177 or inability to phosphorylate T146 was tolerated (Figure 2). Cells harboring T73A and to a lesser extent S177A showed minor growth defects, suggesting phosphorylation of either of these residues could stimulate growth. In contrast, eIF4A phosphomimetic mutations T73D and T146D were unable to support growth, suggesting constitutive phosphorylation of either site prevents cell growth. In each case however, the presence of an additional WT copy of eIF4A (Figure 2, compare plates minus and plus 5-FOA) was sufficient to retain WT growth levels, indicating that mimicking constitutive phosphorylation of these residues did not confer a dominant effect, and suggesting some degree of phosphorylation of these residues is tolerated, so long as protein that can be dephosphorylated is present. In agreement with this result, we were unable to generate a strain harboring the T146D mutation in the *TIF1* and *TIF2* genes in the genome using CRISPR/Cas9 and homology-directed repair with a strategy that was extremely efficient in simultaneously mutating both genes to generate the tolerated T146A mutant (all colonies sequenced had the desired mutations (20)). This further suggests that the presumably lethal effect of mimicking constitutive phosphorylation of T146 is not specific to the *TIF1* paralog of the gene encoding eIF4A. The slow-growth defects of the T73A and S177A mutants combined with the presumed lethality of mimicking constitutive phosphorylation of T73 and T146 suggest dynamic phosphorylation of these residues in eIF4A is needed for optimal growth.

### Mimicking constitutive phosphorylation of two phosphosites arrested cells in mitosis

Because it was previously reported that phosphorylation of *A. thaliana* T164 (equivalent of T146 in yeast) accumulated during mitosis and inhibited growth, we next determined whether the T73D or T146D mutations led to cell cycle arrest in yeast (18). Cells harboring only T73D or T146D mutant copies of the *TIF1* gene did not propagate, so we examined the plated, presumably dead cells following eviction of WT on a 5-FOA plate using differential interference contrast (DIC) microscopy (Figure 3A). We found that WT eIF4A-containing cells appeared asynchronous as expected, with a mixture of cells with and without buds and with some mother-daughter pairs that had not divided. However, cells harboring the phosphomimetic mutations T73D and T146D presented almost entirely as large mother-daughter pairs, suggesting arrest prior to division. This further suggests that activity of eIF4A may be needed for synthesis of a protein required to complete cell division. In contrast, we found that the tolerated phosphodeficient T73A and T146A mutations produced mixtures of cells in different stages similar to WT asynchronous cultures, indicating that preventing phosphorylation of these residues does not arrest the cell cycle. Likewise, both the tolerated S177A and S177D behaved similar to WT, suggesting that S177 phosphorylation, while being diametrically enriched during G1/S arrest, does not arrest the cell cycle, and plays a distinct role from that of T73 and T146 modifications.

**Figure 3.**
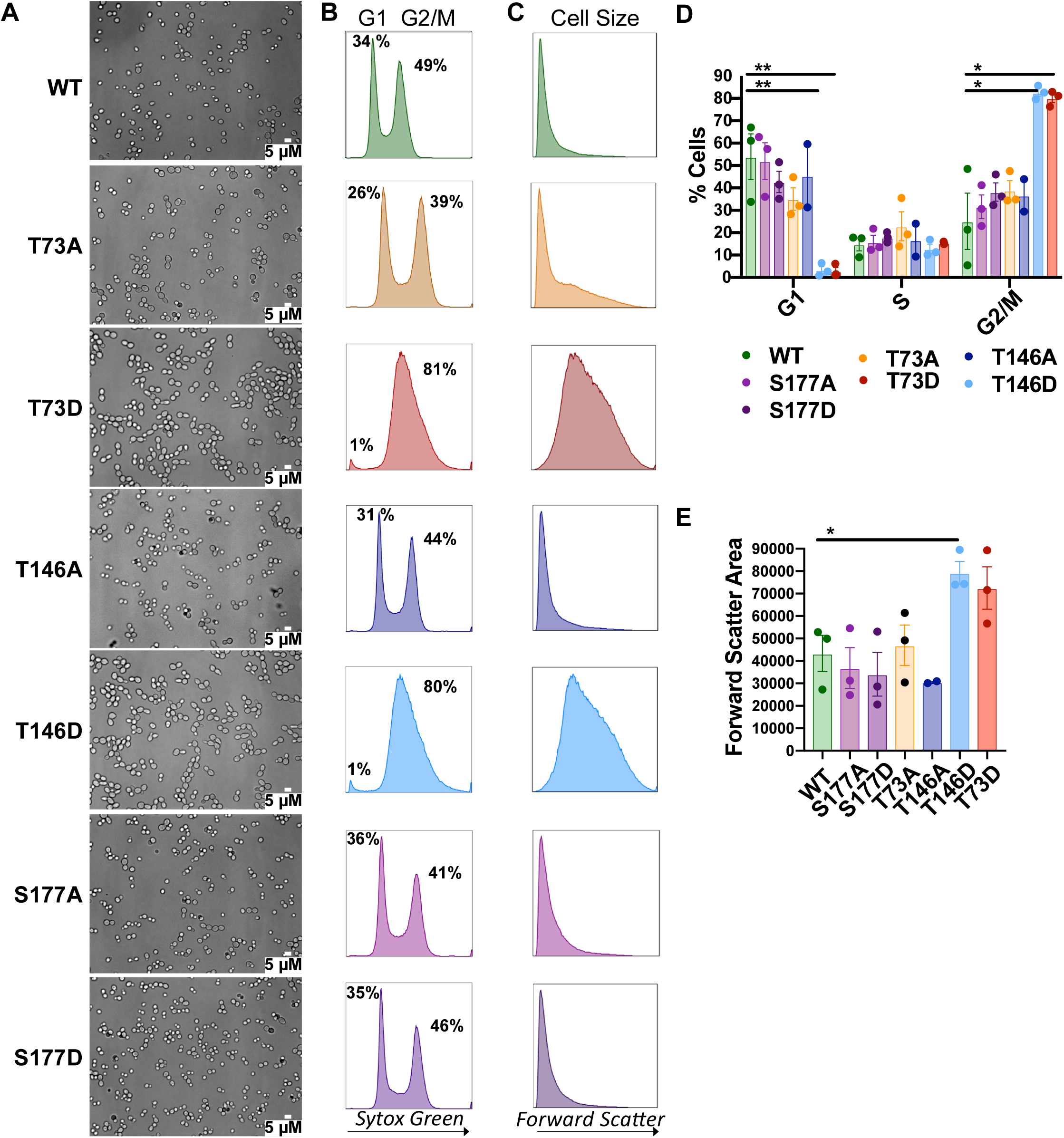
Mimicking constitutive phosphorylation of sites in eIF4A arrests cells in mitosis. A. Yeast cells harboring phosphosite mutations were plated on 5-FOA-containing media to evict the WT copy of the *TIF1*/eIF4A gene and visualized by differential interference contrast microscopy. Cells that did not propagate (T73D- and T146D-only cells) arrested as large mother-daughter pairs. **B, D**. Percentage of individual cells in G1 and G2/M following growth on 5-FOA and fixation and staining by SYTOX green were quantified by flow cytometry. **C, E**. The degree of forward scatter was monitored as an indication of individual cell sizes. Cells that appeared as large-mother daughter pairs by microscopy showed significantly higher percentage of cells in G2/M, with a single broad G2/M peak (**D**, p value <0.01) and significantly higher forward scatter indicative of larger average cell size (**E**, p value <0.05).

To quantitate the effects of these eIF4A mutations on cell division, we assessed DNA content and cell size via SYTOX green staining as previously described (26). Flow cytometric analysis of WT cells showed two peaks indicative of cells with 1N and 2N DNA content (cells in G1 and G2/M, respectively), with a region between the two peaks attributed to cells in S phase. Whereas WT cultures showed peaks consistent with roughly half the cells in G1 or S and half in G2/M, we observed a single broad peak for cells harboring T73D or T146D substitutions of eIF4A, indicating substantially increased percentage of cells within G2/M phases as a function of the two phosphomimetic mutations (81.1% and 79.6%, respectively compared to 48.5% in WT; Figure 3B). In line with these results and the microscopy observations in Figure 3A, the T73D and T146D cells also showed significantly higher values for Forward Scatter (primarily caused by diffraction of light around a cell), indicating larger cell diameter. This suggests that these mutations prevented division of the cells, but not growth of individual cells prior to division (Figure 3C). In contrast, the tolerated mutations T73A, T146A, S177A and S177D did not markedly alter cell cycle stage occupancy or size compared to the WT control (Figure 3A-C).

### RNA binding by eIF4A is inhibited by T73D and T146D

Because eIF4A harboring the T73D and T146D phosphomimetic mutations did not support growth, we were unable to determine the mechanism by which these mutations affect eIF4A function in yeast cells. Instead, we purified recombinant proteins harboring phosphodeficient and phosphomimetic mutations and compared the binding affinity of these variants to that of WT eIF4A for an unstructured RNA in vitro (Figure 4A). We found that consistent with our previous work, titration of unlabeled eIF4A produced a hyperbolic increase in fluorescence anisotropy of a fluorescein-labeled poly(UC) 15-mer, and derived a K_D_ for WT eIF4A of 2.7 ± 0.4 µM (24, 29). Phosphodeficient T73A eIF4A showed similar RNA binding affinity (K_D_ of 2.2 ± 0.4 µM), albeit with a lower overall anisotropy change, suggesting an altered conformation or dynamics of the RNA or protein that do not affect RNA binding affinity. The T146A mutant yielded a marginally higher dissociation constant than WT (5.7 ± 1.2 µM), which is still well below the reported cellular concentration of eIF4A(50-60 µM (30)). However, in keeping with the severe growth defects and cell cycle arrest observed for strains harboring these mutations, no change in anisotropy was observed for phosphomimetic T73D or T146D proteins, at up to 10 µM protein, indicating mimicking constitutive phosphorylation of either residue rendered the protein defective in binding RNA (Figure 4A). In contrast, we found that S177A and S177D did not substantially affect the K_D_ of the protein for RNA (K_D_ of 3.8 ± 0.6 µM and 2.6 ± 0.3 µM for S177A and S177D, respectively.

**Figure 4.**
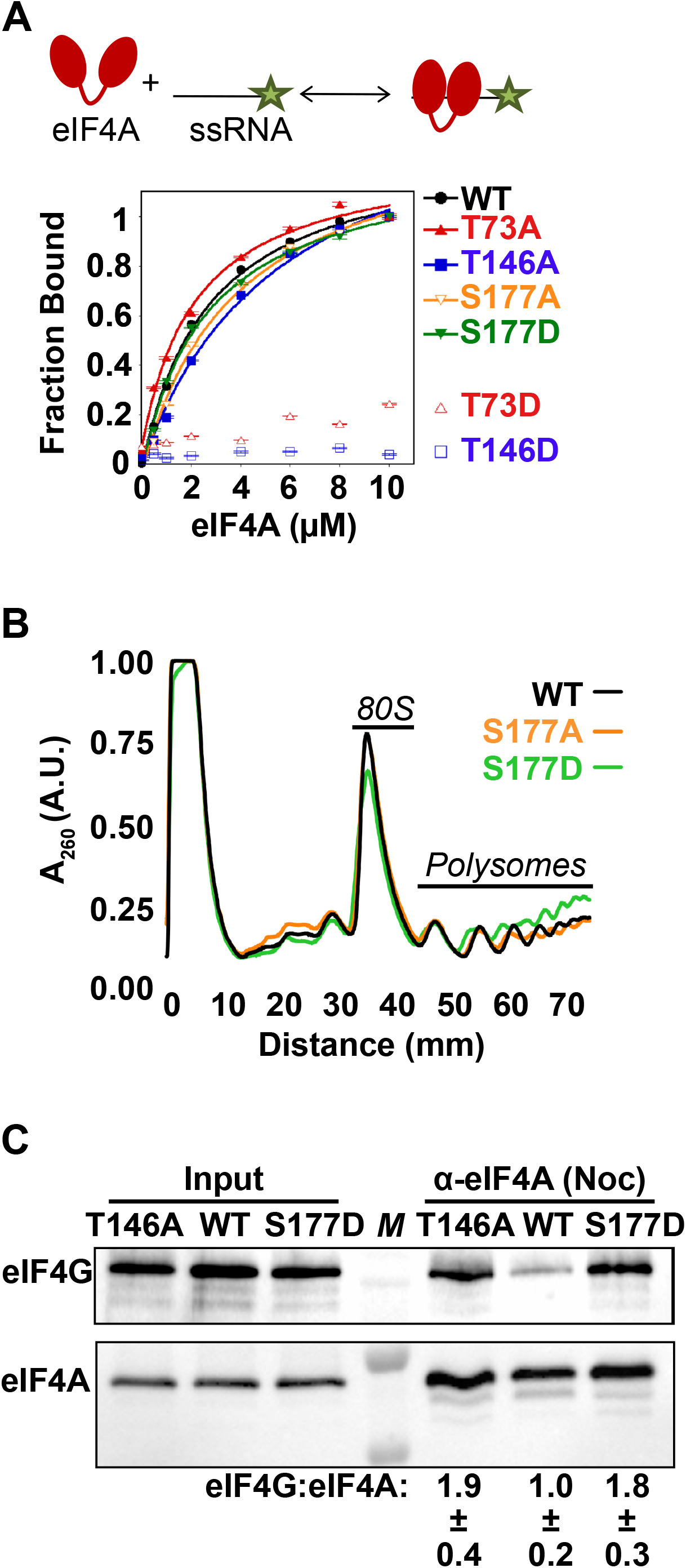
Mimicking phosphorylation of alternative sites near the DEAD box can enhance or repress eIF4A translation activities. **A**. Binding of a fluorescent model unstructured RNA (15 nucleotide poly[UC]) to unlabeled eIF4A with indicated phosphosite substitutions was measured as an increase in fluorescence anisotropy (r). Raw data were converted to fraction bound using the amplitude from r-r_0_ versus concentration plots and fit to a hyperbolic equation to determine K_D_ for WT (black filled circles; 2.7 ± 0.4 µM), T73A (red filled triangles, 2.2 ± 0.4 µM), T146A (blue filled squares, 5.7 ± 0.1 µM), S177A (orange open triangles, 3.8 ± 0.6), or S177D (green filled triangles, 2.6 ± 0.3 µM). The overall anisotropy change was lower for T73A, (amplitude of 0.03 ± 0.01 versus 0.11 ± 0.01 for WT, T146A, S177A, and S177D), suggesting this mutation confers a more restricted conformation of the RNA or alters dynamics of the protein. **B**. Lysates from cells harboring WT and phosphosite-mutated eIF4A expression vectors were fractionated over 10-50% sucrose gradients and monitored by absorbance at 254 nm. P:M ratio, an indicator of global translation initiation rates, was averaged from three biological replicates after calculating the area under the curve of indicated polysome and monosome peaks for cells harboring WT (1.24 ± 0.05), S177A (1.17 ± 0.07), and S177D (1.62 ± 0.10) eIF4A. **C**. The amount of eIF4G co-immunoprecipitation with eIF4A harboring indicated mutations following cell cycle arrest was quantified by western blotting from three biological replicates, and normalized to eIF4A levels, which were consistent across samples.

### T146D was not unfolded, suggesting inhibitory affects are due to mimicking phosphorylation

The T146 residue is surface-exposed, and the phosphorylated version of the residue has been repeatedly detected in peptides in large-scale mass spectrometry studies (16, 31-34), suggesting the effects of phosphomimetic mutations of this residue are likely due to defective RNA binding rather than unfolded protein. However, to assess the latter possibility, we performed circular dichroism spectroscopy on the purified proteins, and then calculated the likely degree of α-helical and β-sheet content using the K2D3 webserver (35)(Figure S1). The calculated α-helical and β-sheet content for the wild-type eIF4A were 37.8 ± 2.2% and 16.8 ± 1.5%, in agreement with predicted values of 39.9% and 18.5% based on sequence. Phosphodeficient T146A was likewise calculated to be comprised of 42.2 ± 2.1% α-helix and 15.1 ± 0.8% β-sheet, e.g., a comparable level to that of WT. Interestingly, the T146D protein was calculated to possess slightly elevated α-helical content (55.3 ± 4.5%) and slightly reduced β-sheets (7.9 ± 1.5%, Figure S1). This could suggest that the phosphorylated protein and its mimic occupy a different conformation or that some regions have been folded differently, but it also indicates that the phosphomimetic protein was not unfolded as a result of the mutation. We also did not see a reduction of tagged-eIF4A as a function of the T146D mutation in a strain expressing untagged WT eIF4A, further suggesting the mutated protein was well-folded (data not shown). Together these results suggest that the surface-exposed T146 residue inhibits eIF4A activity through RNA binding.

### Translation initiation is stimulated by phosphorylation of S177

Because S177 was the major site showing asymmetric differences in phosphorylation as a function of growth phase and mutations of this residue were tolerated, we then opted to determine the effects of the phosphodeficient and phosphomimetic mutations, S177A and S177D, on translation by polysome profiling. Log phase cultures of each strain were treated with cycloheximide, and the lysates then ultracentrifuged on 10-50% sucrose gradients to fractionate RNA by mass, which is then used to globally assess numbers of ribosomes bound per mRNA. The area under polysomes versus monosome peaks determined by A_260_ trace was compared to determine whether the number of ribosomes per mRNA, an indication of cellular translation initiation rates, differed in these strains. We found that S177A cells showed a similar polysome to monosome (P:M) ratio compared to WT, suggestive of similar translation initiation rate when S177 was unable to be phosphorylated (Figure 4B). In contrast, S177D cells showed a slightly higher polysome to monosome ratio, suggesting that mimicking constitutive phosphorylation of S177 enhanced translation initiation rate. Together these results suggest that phosphorylation of S177 promotes translation initiation, consistent with its accumulation during G1/S and absence during G2/M arrest, when translation is repressed (27).

### Mimicking constitutive phosphorylation of S177 or preventing phosphorylation of T146 of eIF4A enhanced eIF4G interaction during mitosis

We then analyzed the effects of preventing changes in phosphorylation observed to occur during mitosis by either mimicking constitutive S177 phosphorylation (S177D) or preventing T146 phosphorylation (T146A) during nocodazole arrest. We immunoprecipitated WT eIF4A or eIF4A harboring T146A or S177D from cells after 2 hours treatment with nocodazole, and blotted the samples for eIF4G (Figure 4C). Interestingly, we found that either substitution led to enhanced co-immunoprecipitation of eIF4G with eIF4A compared to WT eIF4A. This suggests that both phosphorylation of T146 and dephosphorylation of S177 could contribute to diminished eIF4A•eIF4G interaction observed during mitotic arrest (Figure 1A), and highlighting the role of these events in normal mitotic translation behavior.

## Discussion

The eukaryotic translation initiation factor eIF4A plays a critical role in remodeling mRNA•protein complexes (mRNPs) and stimulating translation, yet only one posttranslational modification of this protein has been studied in detail (18). Here we analyzed cell-cycle-dependent phosphorylation of eIF4A as a means of translation regulation in yeast. We found that eIF4A harbors three surface-exposed phosphorylation sites proximal to the DEAD-box motif that show changes in phosphorylation capable of regulating function during cell cycle transitions. Importantly, these sites are conserved across eukarya, suggesting they could be universal inputs for both enhancement and rapid shutdown of translation and other activities of eIF4A.

First, immunoprecipitation-mass spectrometry experiments revealed the highest degree of phosphorylation was observed for residue S177, which was completely phosphorylated in G1/S and dephosphorylated in G2/M arrest, suggesting this is the major phosphorylation event regulating eIF4A activity between G1/S and G2/M in yeast. Phosphorylation of this site is likely to stimulate eIF4A activity as mimicking constitutive phosphorylation led to enhanced interaction with eIF4G during G2/M arrest and an increase in polysome:monosome formation, indicative of increased translation initiation rate. In contrast, a phosphodeficient mutant decreased polysomes and a modest growth defect. The eIF4G binding site is on the opposite face of eIF4A from its DEAD box motif, which is immediately adjacent to S177. This suggests S177 is not likely to directly interact with eIF4G, but instead suggests that S177 phosphorylation stimulates the transition to or stabilizes the closed state of eIF4A, which harbors high affinity for eIF4G and RNA.

In contrast, roughly equivalent levels of phosphorylation at T73 and T146 were observed during G1/S or G2/M, respectively. Mimicking constitutive phosphorylation of either of these sites arrested cells as large mother-daughter pairs prior to division in a state similar to Mitotic arrest, and prevented eIF4A•RNA binding in vitro, indicating phosphorylation of these sites inhibits eIF4A activity. Moreover, the fact that inhibiting eIF4A activity through these phosphomimetic mutations prevented division suggests that eIF4A activity is needed for completing cell division. The inability of either phosphomimetic mutant to bind RNA suggests phosphorylation of these two sites impedes RNA binding through the placement of a negatively charged phosphate that directly opposes the RNA backbone, and/or by promoting an open, low RNA- and eIF4G-affinity conformation of eIF4A. In agreement with the latter possibility, we observed rescue of eIF4G association with the T146A protein during mitosis, when eIF4A•eIF4G interaction is normally diminished. This suggests that phosphorylation of eIF4A at T146 during mitosis is capable of decreasing that interaction.

Because T73 and T146 showed roughly the same extent of phosphorylation (up to 24% and 27% phosphorylated peptides, respectively), but T73 phosphorylation increased in G1/S and T146 phosphorylation accumulated during G2/M arrest, it is likely that these two events could offset in the different cell cycle phases. Taken together with the modest degree of change in phosphorylation of these residues, these observations suggest that T73 and T146 phosphorylation may normally serve to respond to other potentially more acute stressors, such as glucose deprivation or heat shock, both of which result in rapid eIF4A release from mRNA in yeast (36). It will be of interest to follow up on whether phosphorylation of these sites play a role in such stress responses in future work.

Instead, our data suggest that phosphorylation of S177 plays the principal role in enhancing eIF4A activity during G1/S in yeast (Figure 5). Because we also observed that blocking eIF4A activity via T146D or T73D mutations prevented cells from completing the mitosis to G1 transition (Figure 3), it is likely that activity of eIF4A is needed for the translation of one or multiple transcripts required for completing cell division. Phosphorylation of S177 may enhance this activity, and dephosphorylation may dampen eIF4A activity prior to that transition to ensure a transcript is not prematurely translated. Previous work in yeast showed that proper control of expression of the G1 cyclin Cln3 requires a repressive upstream open reading frame in the 5’UTR of the *CLN3* mRNA (37). Deletion of that uORF led to decoupling of growth and division, suggesting repression of Cln3 synthesis in other phases of the cell cycle prevents premature transition to G1. It is possible that dampened eIF4A activity during mitosis and enhancement upon reentry into G1, mediated by the changes in phosphorylation observed here, plays a role in the proper timing of the *CLN3* ORF translation. Alternatively, eIF4A phosphorylation changes could influence the timing of translation of other cell cycle transcripts such as cdc25 mRNAs, noted as eIF4A-dependent transcripts in fission yeast that influence entry into mitosis (38). Recent work in mammalian cells reported that both translation repression and mRNA turnover occur during mitosis (27, 39). In particular, work by Tanenbaum et al indicated that repression of a transcript for an inhibitor of the anaphase promoting complex occurs in mammalian cells, which then allows cells to complete the mitosis to G1 transition (27). It is possible that inhibition of eIF4A activity by dephosphorylation of S177, or phosphorylation of T146 could play a role in repressing transcripts in yeast that serve a similar purpose.

**Figure 5.**
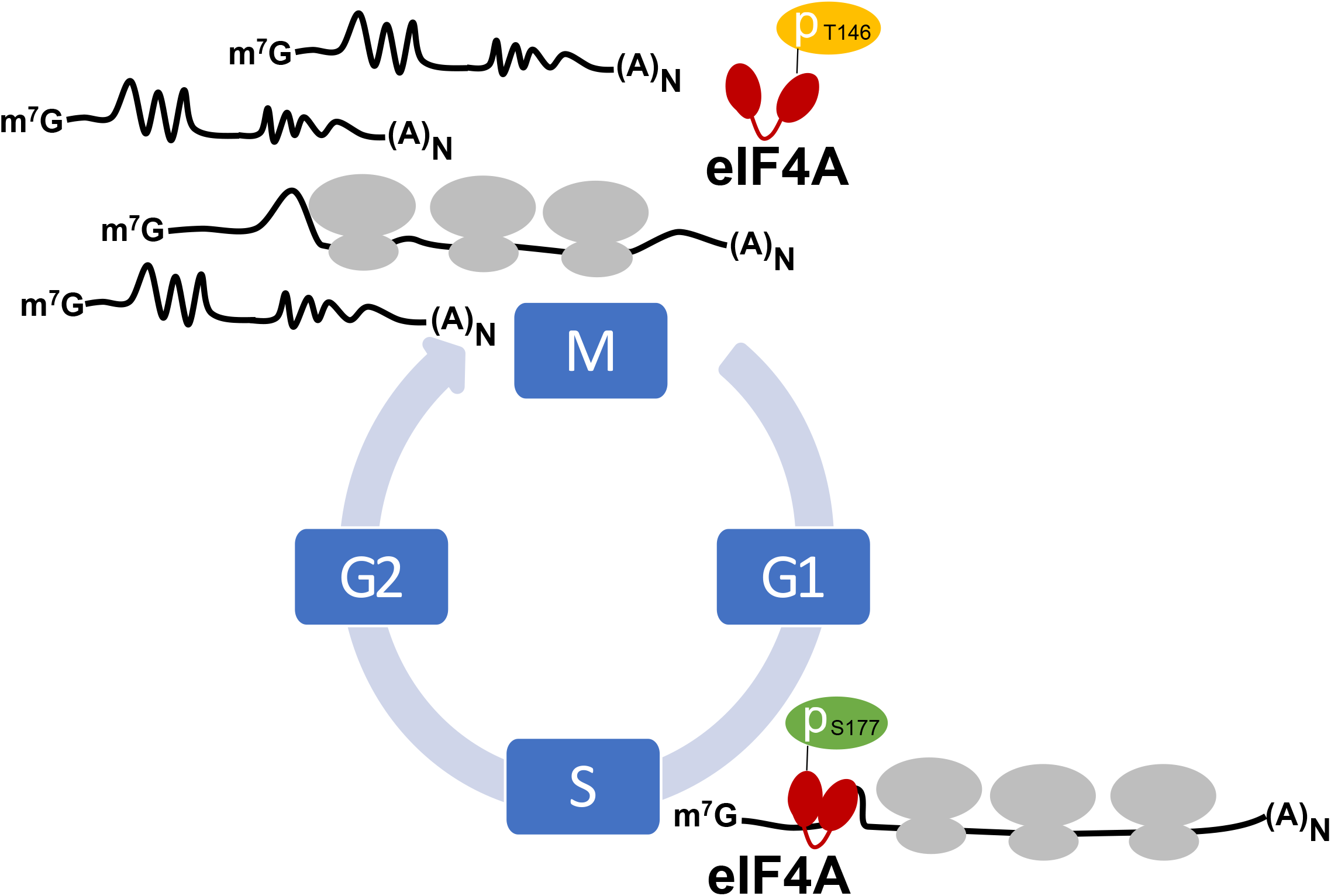
Model for regulation of eIF4A activity during the cell cycle. Phosphorylation of T146 and dephosphoryation of S177 residues together decrease eIF4A activities leading to reduced ribosome loading and translation of eIF4A-dependent mRNAs during mitosis. The inverse effects are predicted to promote translation of an alternative group of transcripts and allow completion of cell division and subsequent growth during G1 and S phase.

How might eIF4A inhibition affect these translation events during the cell cycle? It has been demonstrated that certain transcripts exhibit more and less dependence on eIF4A for translation (4, 40, 41), and that translational control is intimately linked to the cell cycle (42). It is possible that limiting eIF4A activity during mitosis could lead to translation of a different group of messages. These mRNAs could harbor less structure, or sequence/structure motifs that make them more competitive for eIF4A, eIF4G, or ribosome binding (41, 43-46). In fact, previous studies suggested that cap-independent translation mechanisms could be at play during mitosis (45, 46). Inhibiting eIF4A activity through the phosphorylation changes described here would be expected to dampen the ability of most transcripts to associate with ribosomes, making transcripts that harbor eIF4A- or other cap-independent translation elements better able to compete for ribosome binding, in agreement with both studies suggesting cap-independent translation during mitosis, as well as the recent report that translation decreases during mitosis are more prevalent (27, 45, 46). Further analysis of the translational outcomes of modulating eIF4A activity through phosphorylation changes during mitosis will be of great interest in future work, and may hold relevance for inhibition of mitosis in cancer cells or treatment of pathogenic yeast.

## Acknowledgements

The authors would like to thank Austin D’Angelo, Paul Cullen, Paul Gollnick, Qian He, Xiaozhuo Liu, Laura Rusche, and Michael Yu for providing strains, reagents, and helpful suggestions for this project. This work was supported by a Mark Diamond Research Fund Fellowship from the University at Buffalo (A.S.), NIH grants R00GM119173 and R01GM139977 (S.E.W.), and institutional funds from the University at Buffalo College of Arts and Sciences (S.E.W.) and Roswell Park Comprehensive Cancer Center (J.B.). This work involved the Flow and Image Cytometry Shared Resource at Roswell Park supported by National Cancer Institute (NCI) grant P30-CA016056.

